# Distinct survival, growth lag, and ribosomal RNA degradation kinetics during long-term starvation for carbon or phosphate

**DOI:** 10.1101/2020.09.04.284034

**Authors:** Yusuke Himeoka, Bertil Gummesson, Michael A. Sørensen, Sine Lo Svenningsen, Namiko Mitarai

## Abstract

Stationary phase is the general term for the state a bacterial culture reaches when no further increase in cell number occurs due to the exhaustion of nutrients in the growth medium. Depending on the type of nutrient that is first depleted, the metabolic state of the stationary phase cells may vary greatly, and the subsistence strategies that best support cell survival may differ. As ribosomes play a central role in bacterial growth and energy expenditure, ribosome preservation is a key element of such strategies. To investigate the degree of ribosome preservation during long-term starvation, we compared the dynamics of ribosomal RNA (rRNA) levels of carbon-starved and phosphorus-starved *Escherichia coli* cultures for up to 28 days. The starved cultures’ contents of full-length 16S and 23S rRNA decreased exponentially and phosphorus starvation resulted in much more rapid rRNA degradation than carbon starvation. Bacterial survival kinetics were also quantified over the starvation period. Upon replenishment of the nutrient in question, carbon-starved cells resumed growth faster than cells starved for phosphate for the equivalent amount of time, and for both conditions, the lag time increased with the starvation time. While these results are in accordance with the hypothesis that cells with a larger ribosome pool recover more readily upon replenishment of nutrients, we also observed that the lag time kept increasing with increasing starvation time, also when the amount of rRNA per viable cell remained constant.

**Importance:** Bacteria grow exponentially consuming nutrients, and then starve until the next nutrient is added. To elucidate the survival kinetics of the cells under starvation, we performed month-long, carbon and phosphorus starvation experiments of *Escherichia coli* monitoring ribosomal RNA levels and survival of the cells. The starved cultures’ concentration of ribosomal RNA dropped exponentially with time, and the speed of degradation was much quicker under the phosphorus starvation than the carbon starvation. We have also quantified the lag time, i.e., the time needed to resume growth when the starved cells are transferred into fresh media. The observation revealed that the lag time increases with starvation time and the phosphorus starvation has a greater impact on the increase of the lag time.

## Introduction

The amount of protein production machinery in a bacterial cell is tightly regulated during exponential growth, which is important for the proper allocation of the available resources to maximize protein synthesis and thereby growth rate (Maaløe 1979, Scott et al. 2011). The increased demand for protein synthesis capacity at higher growth rates can be appreciated as a linear increase in the concentration of ribosomes (estimated by the RNA/protein ratio) as a function of the steady-state exponential growth rate (Schaechter et al 1958; Scott et al. 2010). This concept was recently extended to consider relatively slow exponential growth rates obtained in chemostats by limiting a specific component of the growth medium (Li et al. 2018). There, it was found that the RNA/protein ratio is about 2-fold lower under phosphorus-limited growth compared to the carbon-limited case for a given growth rate. Since more than 60% of cellular phosphate is found in ribosomal RNA (rRNA), it is not surprising that cells would have evolved mechanisms to reduce ribosomes to a minimal number during phosphorus-limited growth conditions. This finding also indicates that cells are sustaining “unnecessary” ribosomes in the carbon-limited case, which was confirmed by an accumulation of mRNA-free ribosomes under this growth condition (Li et al. 2018). An overcapacity of ribosomes has been noted previously, and was proposed to enable a rapid response to a nutrient upshift (Kock 1971, Mori 2017).

Upon severe starvation where the increase of biomass comes to a halt, ribosomes are known to be subject to degradation. This has been demonstrated in various conditions including carbon starvation (Jacobson and Gillespie 1968; Kaplan and Apirion 1975; Mandelstam and Halvorson 1960; Zundel et al. 2009, Fessler 2020) and phosphorus starvation (Julien et al. 1967; Maruyama and Mizuno 1970, Fessler 2020). In a comparative short-term study, we showed that 80 minutes after abrupt starvation introduced by filtration of bacteria into growth medium lacking a particular nutrient type, 10% of rRNA was degraded for carbon starvation, whereas ∼40% was degraded during phosphate starvation (Fessler 2020). In qualitative agreement with this observation, others have reported that 12-24% of ribosomes were degraded after 3-4hs of carbon starvation (Mandelstam and Halvorson 1960; Zundel et al. 2009) and about 60% was degraded after 24h (Zundel et al. 2009), while for phosphorus starvation ribosome degradation was more severe, resulting in 50%-80% degradation after 12h of starvation (Maruyama and Mizuno 1970).

Recent studies of carbon starvation report an exponential decrease in cell viability during several weeks of starvation in minimal medium (Phaiboun et al. 2015; Schink et al. 2019). However, the potential effects of cellular ribosome content on the physiological behavior of the cells, such as their viability and the lag time, i.e., the time required for starved cells to commence growing after the addition of fresh medium, has not been explored under long-term starvation. In this paper, we studied the effects of long-term starvation (up to 28 days) for carbon and phosphorus on ribosome levels, survival kinetics and growth lag time.

## Material and Methods

### Bacterial strain and Media

The wild-type strain *Escherichia coli* K-12 MAS1081 (MG1655 *rph*^+^ *gatC*^+^ *glpR*^+^) was used in all growth experiments. Cultures were kept at 37 °C with shaking at 190-220 rpm in a water bath, in morpholinepropanesulfonic acid (MOPS) minimal medium supplemented with 0.2 % glucose (Neidhardt et al 1974). Cell growth was monitored spectrophotometrically by optical density at 436 nm (OD_436_) and cultures were grown exponentially for at least fifteen generations before being exposed to starvation. Carbon or phosphorus starvation was introduced gradually by transferring cells from balanced growth cultures to preheated MOPS medium supplemented with 0.07% glucose (designated here as MOPS low carbon (LC) medium) or MOPS medium with 0.2% glucose but with reduced dipotassium phosphate (0.066mM instead of 1.32mM) (designated here as MOPS low phosphate (LP) medium), and letting the cultures enter stationary phase due to the lack of carbon source (LC cultures) or phosphorus source (LP cultures).

For RNA measurements by Northern blot, whole-cell spike-in cultures of the *E. coli* strain MAS1074 were used for normalization. MAS1074 expresses the *selC* gene encoding tRNA^selC^ under the control of an isopropyl β-D-1-thiogalactopyranoside (IPTG)-inducible promoter (Svenningsen et al 2017). The spike-in strain was grown in MOPS medium with 0.2% glucose and 100 μg/ml ampicillin at 37 °C. tRNA^selC^ expression was induced at OD_436_ 0.10 by adding IPTG to a final concentration of 1mM. After 4 hours of inducing conditions (at OD_436_ 0.42), the spike-in culture was aliquoted in a 1:2 ratio with RNA*later*™ solution (Invitrogen) on ice and stored at -80 °C.

### Viability measurement

The viability was determined by counting the number of colony-forming units (CFU) on Luria-Bertani (LB) agar plates. After plating, the plates were incubated at 37°C. The number of colonies was counted after one overnight incubation as well as after 2 days of incubation. The CFU did not change after 2 days of incubation as assessed by counting the plates again after 3 days of incubation and obtaining identical numbers. Serial dilutions were performed so that the number of colonies was 100∼200 per plate, while it occasionally dropped below 50 especially in the phosphorus-starvation cultures. The average values and standard deviations reported for viability measurements were based on at least four plates.

### Spike-In Normalisation and RNA Extraction

For rRNA measurements, bacterial culture samples were harvested by mixing an aliquot of the culture with 1/5 vol of a stop-solution composed of 5% water-saturated phenol in ethanol at 0°C (Bernstein et al 2002). Prior to total RNA extraction, a volume of spike-in culture corresponding to roughly 5% of the experimental sample was added (the sample and spike-in volumes are listed in Supplementary Data 1). Total RNA was extracted by hot phenol as described in Gummesson et al 2020.

### Northern Blots and rRNA quantification

Northern blots were performed as described in Gummesson et al 2020. 5’end-labeled oligo-DNA probes (γ-[32P]-ATP; PerkinElmer) complementary to a sequence in the 16S rRNA, 23S rRNA or SelC tRNA were used to detect the immobilized RNA on the membranes (probe sequences are listed in Supplementary Text Table.S1). The radioactivity present in specific bands was measured on a Typhoon phosphor Imager FLA7000 (GE Healthcare) at 100 microns, and quantified usingImageQuant TL 8.2. The quantified intensity of each rRNA band was first adjusted to account for the contribution from the spike-in cells using a lane on the same Northern blot that was loaded with spike-in cells only. Then, the corrected rRNA signal intensity was divided with the SelC tRNA signal in the same lane to give the normalized rRNA level. For Fig. 1B, this ratio is plotted relative to the average of the two samples harvested on day 0, while for Fig. 1E+1F, the ratio is plotted relative to the average of measurements from the culture prior to starvation. While changes in OD provide a clear measure of the growth of bacterial cultures in the steady state, where all cell constituents grow with the same rate, changes in optical density are complex to interpret upon disruption of the steady state, e.g. by starvation (Fessler et al 2020, Koch 1970). Accordingly, the correlation between OD measurements and bacterial concentration (CFU/ml) was good for exponentially growing cultures, but poor for the starved cultures (Supplementary Fig.S1). Therefore, measurements of rRNA levels from starved cultures were retroactively normalized to the spike-in signal from SelC tRNA only relative to the culture sample volume, disregarding changes to the OD that had occurred after the entry to stationary phase.

**FIG 1:**
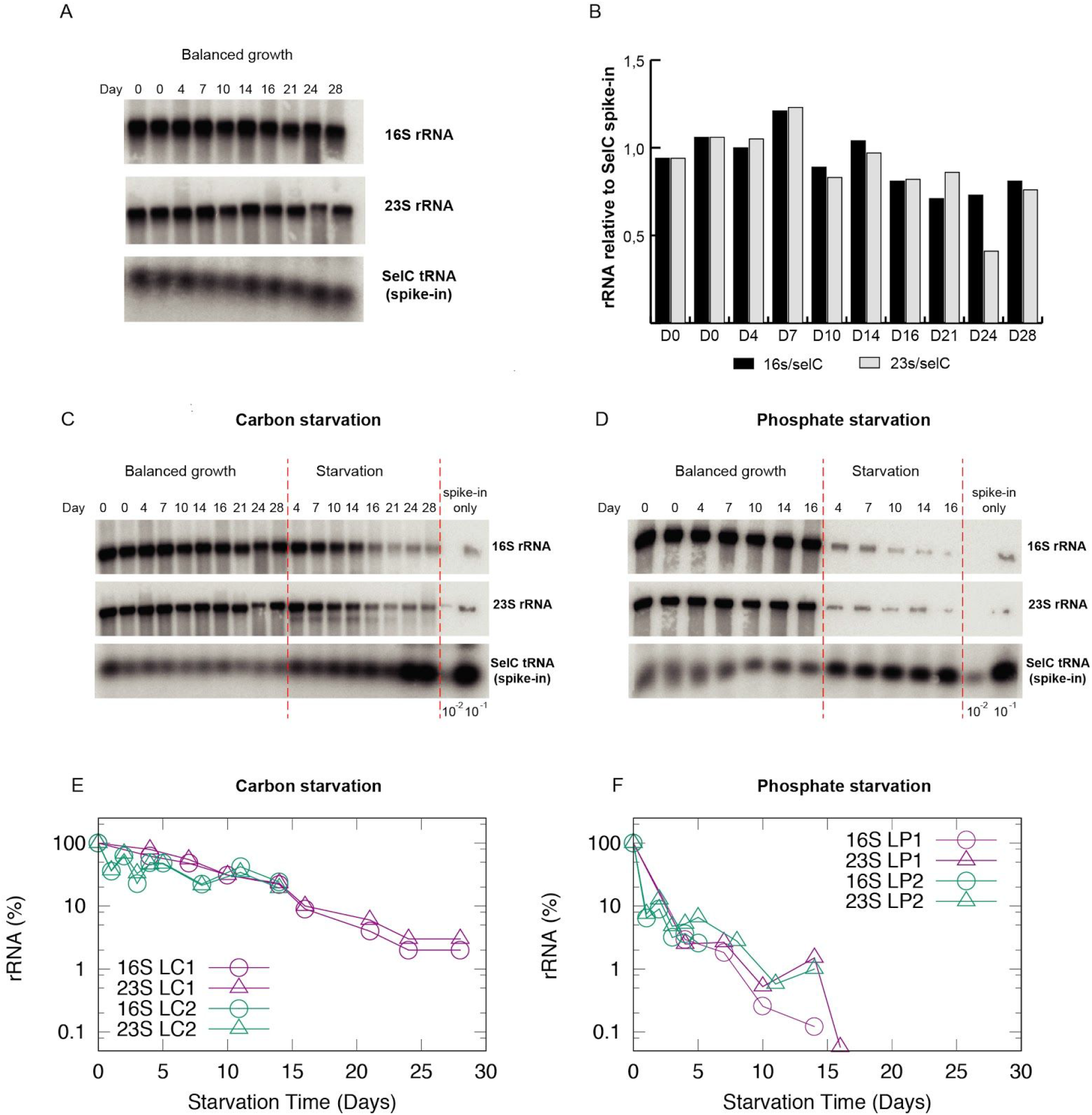
Quantification of ribosomal RNA in cultures undergoing long-term starvation. (A) Northern blot of total RNA from samples harvested during balanced growth. Day 0 indicates the start of the experiment and Day 4, 7, 10, 14, 16, 21, 24 and 28 indicates the number of days the spike-in cells were stored prior to mixing with a fresh aliquot of cells in balanced growth. The resulting blot was probed for 23S, 16S and the spike-in-cell-specific tRNA SelC as indicated. (B) The levels of rRNA (16S; black bars, 23S; grey bars) were quantified by normalizing to tRNA SelC from the spike-in cells and shown relative to the average of the two RNA samples harvested on Day 0. (C) Northern blot of total RNA from cultures starved for carbon (middle panel, Day 4, 7, 10, 14, 16, 21, 24 and 28 indicates the number of starvation days) as well as control cultures in balanced growth prepared fresh on the same days (left panel). (D) Northern blot of total RNA from cultures starved for phosphate (middle panel, Day 4, 7, 10, 14, and 16 indicates the number of starvation days) as well as control cultures in balanced growth prepared fresh on the same days (left panel). For both Northern blots in (C) and (D), the right panel shows 1/100 and 1/10 dilutions of samples harvested from the spike-in cells. The blots were probed for 23S rRNA, 16S rRNA and tRNA SelC as indicated. (E) The plots show 16S and 23S rRNA levels relative to balanced growth during carbon starvation and (F) shows the quantification of 16S and 23S rRNA during phosphate starvation. Independent measurements are labeled as LC1 and LC2 for carbon starvation, and as LP1, LP2 for phosphate starvation.

### Lag time measurements

Each measurement day, an aliquot of the starved culture was diluted into MOPS minimal medium. 200µl of the diluted culture was placed in each well of a microtiter plate (TPP 96-well flat bottom, or greiner 96-well flat bottom), and the optical density (*OD*_436_) was measured in a microtiter plate reader with shaking at 37°C (FLUOstar OMEGA, BMG LABTECH) every 5 minutes for about 24 hours. The culture aliquots were diluted by fresh medium so that the OD_436_ of the diluted culture was above the detection limit of the plate reader which we determined as ∼ 0.05. The cultures were typically diluted by 10∼20 fold. Four specific wells of the microtiter plates formed droplets on the lid repeatedly and gave inconsistent OD readings. The corresponding four OD curves were therefore removed from the data sets, and average growth curves from the remaining 84 wells were computed.

### Fitting of the growth curves

The obtained time courses of the optical density were fitted by a combination of constant line f and a simple exponential curve g to determine the growth rate in exponential phase and the duration of the lag time in each well. The fitting regions for the lag phase (by a constant line) and exponential phase (by an exponential curve) were determined manually. The fittings were discarded if the average square difference between the fitting line and data,

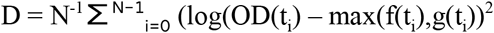

exceeded 0.1, where t_i_ is the time after at the ith measurement of OD curve, and t_N-1_ is the upper bound of the fitting region for the exponential phase. The lag time of each well is estimated by the fitting as the cross point of f and g. The apparent lag time is given as the average of the lag times of all the wells. The standard error is computed by propagating the unbiased standard deviations of *f* and *g*. Calculations for subtracting the death effect from the lag time are explained in section 2 in Supplementary Text.

## Results

### Quantification of rRNA in long-term starved *E. coli* cells

Degradation of rRNA during starvation for hours to a few days has been reported by us and others (Jacobson and Gillespie 1968; Kaplan and Apirion 1975; Mandelstam and Halvorson 1960; Zundel et al. 2009, Fessler 2020). In order to accurately determine the relative rRNA levels over the long-term starvation period examined here, it was necessary to first establish a quantification method that allowed a meaningful comparison of relative RNA levels in culture aliquots harvested on different days. Since the efficiency of RNA extraction can vary from sample to sample, it is necessary to normalize the measured levels of the RNA of interest to a suitable reference RNA, but given the month-long duration of the starvation period, no endogenous RNAs could be expected to remain at the same level throughout the experiment. A whole-cell spike-in method is useful for such circumstances (Gummesson et al. 2020) and is carried out by spiking each experimental sample with a small fixed relative volume of reference bacterial cells, followed by normalization of the level of the RNA of interest to one or more RNA species that are highly abundant in the spike-in cells and absent in the experimental samples. We used a spike-in culture of *E. coli* that overexpresses the rare transfer RNA tRNA^selC^, which is undetectable in the experimental samples (Svenningsen et al 2017, Stenum et al 2017). Aliquots of a spike-in culture were stored in RNAlater, and for each RNA harvest day, a frozen aliquot was defrosted, mixed with the sample culture, and relative rRNA levels were reported as the ratio of rRNA to tRNA^selC^.

To verify that the frozen stocks of spike-in cells carrying tRNA^selC^ could be used for normalization during a long-term experiment, we took advantage of the fact that ribosome content is linearly related to growth rate under steady-state exponential growth (Schaechter et al 1958; Scott et al. 2010) and thus, the rRNA per OD unit of cells should be the same in *E. coli* cultures growing at the same steady-state growth rate, even if measured on separate cultures on different days. To test the method over the course of 28 days, we grew a culture of wildtype *E. coli* K-12 every few days to balanced growth with a doubling time of around 54 min. Culture aliquots were mixed with aliquots of the stored spike-in culture at a fixed ratio of 0.05 spike-in culture to experimental culture, based on OD units. RNA was purified from the cell mixture, stored, and at the end of the 28-day period, we ran a Northern blot of all the samples to test whether the ratio of rRNA to tRNA^selC^ was constant. Fig. 1A shows an example of such a Northern blot, and Fig. 1B shows that the rRNA/spike-in ratio was indeed fairly constant over the 28-day period, varying with a standard deviation of 16% for 16S rRNA/ tRNA^selC^ and 23% for 23S rRNA/tRNA^selC^. The additional variation in 23S rRNA is caused by the sample from day 24, where extensive degradation of 23S rRNA in the sample is noticeable on the Northern blot. No systematic trends (for example, an increase of rRNA over time due to the degradation of *selC* in the frozen stock) could be observed. Thus, a frozen whole-cell spike-in culture allows quantification of relative rRNA levels over at least a 28-day period.

To monitor the long-term effects of starvation for carbon or phosphorus we grew cultures to balanced growth, after which the bacteria were subjected to starvation for the carbon source (glucose) or phosphorus source (potassium diphosphate) by transfer to MOPS low carbon (LC) or MOPS low phosphorus (LP) medium. The cultures remained in the shaking water bath for up to 28 days and were sampled for measurements of rRNA content, cell viability, and growth lag time as described in detail in the Methods section. In both low nutrient media, the cultures reached an OD_436_ of 1-2 within 8-10 hours, and then entered the stationary phase due to the lack of the limiting nutrient.

To determine the rRNA levels in the starved cultures, culture aliquots were mixed with known quantities of the frozen stock spike-in as described for cultures in balanced growth above. Examples of the resulting northern blots are presented in Fig. 1C and 1D, and the results of two independent measurements are shown in Fig. 1E and 1F for the carbon and phosphate starvation, respectively.

In the carbon starvation case (Fig. 1E), the levels of rRNA per culture volume did not decrease detectably for the first several days of starvation but decreased to 50% of the value during balanced growth after approximately two weeks. In the final two weeks, the rRNA concentration decreased monotonically by another 50%. By contrast, rapid and extensive rRNA degradation was observed from the onset of phosphorus-starvation (Fig. 1F). Within the first day of starvation, the rRNA concentration had dropped below 40%, and the decrease continued approximately exponentially with a decay rate of 0.35 per day, dropping below the level at which we could reliably measure it on the Northern blots after 10 days. One limitation of our whole-cell spike-in approach is that the rRNA from the spike-in cells, which contributes insignificantly to the total rRNA when mixed into growing cultures at a 1:20 ratio, overwhelms the rRNA contribution from the starved cultures once a large percentage of their rRNA has been degraded. Relative rRNA levels below ∼5% are therefore too close to the detection limit of our method and are simply evidence that the rRNA levels are “very low” at the qualitative level. In conclusion, cultures that enter the stationary phase due to the lack of phosphate, an integral part of the RNA backbone, show much more extensive rRNA degradation than cultures starved for the carbon and energy source glucose, even several days or weeks into starvation.

### Distinct survival kinetics of *E. coli* during carbon and phosphorus starvation

We wondered how the distinct rRNA degradation dynamics related to cell survival during starvation for the two different types of nutrients. The kinetics of cell death under starvation was studied by measuring the ability of cells from the starved cultures to form colonies within 2 days on rich medium agar plates. The number of viable cells (colony forming units, CFU) per culture volume is depicted in Fig. 2A for the carbon-starved case and 2b for the phosphorus-starvedcase as a function of the time the cells had remained under starvation conditions prior to plating on rich medium. In both cases, viability remained somewhat constant for the first five days of starvation. After that, carbon and phosphorous starvation resulted in remarkably different survival kinetics. The concentration of viable cells in carbon-starved cultures showed a continuous exponential decay, consistent with recent results of carbon-starvation by glycerol depletion (Schink et al., 2019, Biselli et al. 2020), with a decay rate of approximately 0.22 per day. On the other hand, cell viability showed a rapid ∼12-fold drop around day 5 to 6 and then remained unchanged during phosphorus-starvation, except for one replicate (labeled as “LP1” in Fig. 2B), which appeared to show a more continuous decrease in viability. The behavior of the phosphorus-starved cultures is similar to that reported for *E. coli* outgrowth from complex LB medium, containing a stationary phase, and a death phase, followed by a long-term stationary phase (Finkel, 2006).

**FIG 2.**
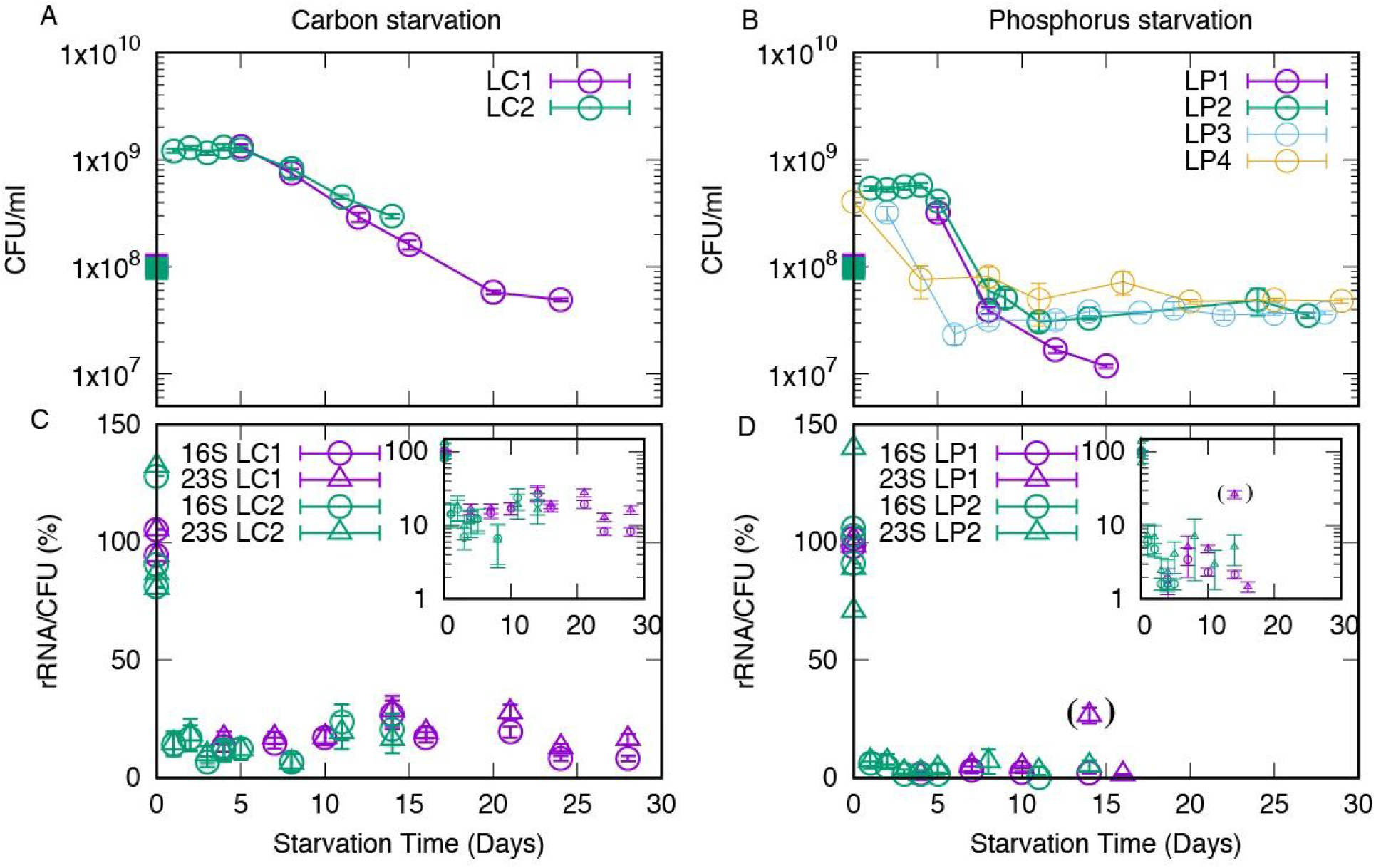
Survival kinetics of *E. coli* during long-term starvation and the population-averaged ribosomal RNA content. Time course of CFU/ml for (A) the carbon starvation case and (B) the phosphorus starvation case. Independent measurements are labeled as LC1 and LC2 for carbon starvation, and as LP1, LP2, LP3, LP4 for phosphate starvation. Square symbols on day 0 show the estimated CFU/ml at the time of RNA harvest from exponentially growing cultures, calculated based on OD_436_. Error bars indicate the unbiased standard error of the mean (N≧3). (C) and (D) show the relative levels of 16S and 23S rRNA per surviving (colony-forming) bacterium. The outlier point in 16S LP1 is disconnected and enclosed in the parenthesis because the value is likely due to experimental error. Error bars indicate the unbiased standard error which is computed by propagating only the unbiased standard deviation of CFU/ml (for more details, see section 1 of Supplementary Text).

To illustrate the average decrease in ribosome levels per viable cell, the percentage of rRNA/CFU relative to cultures in balanced growth are shown in Fig. 2C and 2D. In the case of carbon starvation, the rRNA/CFU ratio stays relatively constant around ∼20% of the level in balanced growth throughout the starvation period (Fig. 2C). The magnitude of the initial drop is at least partly caused by the increase in CFU that occurs without a concomitant increase in biomass when cells undergo reductive division upon entry to stationary phase (Kaprelyants et. al. 1993, Nyström et. al. 2004, Arias et. al. 2012, Gray et. al, 2019). Since reductive cell divisions result in a reduction of the average cell volume, the actual drop in the average intracellular concentration of ribosomes may be 2-4 fold less than the relative rRNA levels depicted in Fig. 2C would suggest.

In the phosphate starvation case, the drop of rRNA/CFU (Fig. 2D) early on is significantly more rapid and severe than the carbon starvation case, showing a 10-fold decrease already after a day, and decreasing to a few percent of the original value by day 4. Again, the fluctuations seen far into phosphate starvation are likely to be explained by imprecise measures of the very low levels of rRNA relative to the contribution from the spike-in cells. For the same reason, we did not attempt to measure rRNA levels past the two-week time point under this condition. Note that the decrease in rRNA levels in the cultures is exponential while cell viability drops step-wise for phosphorus starvation. This results in a trend of increased rRNA/CFU after day 5 (Fig. 2B LP1 and LP2). We suspect this trend might reflect that some full-length rRNA molecules from nonviable cells in the culture are detected on the northern blots.

The finding that phosphorus-starved cells on average survive with much fewer ribosomes per viable cell than carbon-starved cells suggests that more ribosomes than the necessary minimum are maintained in carbon-starved cells also during long-term starvation.

### Growth lag time after the stationary phase depends on the type of starvation

The time interval from starved cells are placed under growth-permitting conditions and until growth is actually observed; the lag time, is a measure of the physiological readiness of the cells to resume active conversion of nutrients into bacterial biomass. Several reports suggest that lag times generally increase with the duration of the starvation time (Augustin et al. 2000; Oscar 2005; Pin and Baranyi 2008, Levin-Reisman 2010) although a unifying molecular mechanism underlying this correlation has not been identified. Since protein synthesis activity by the ribosomes is a prerequisite for cell growth, one could naively expect that a progressive loss of ribosomes during starvation could be the underlying cause of the positive correlation between starvation time and the subsequent growth lag time.

We measured the lag time of populations of *E. coli* as a function of starvation time in medium lacking carbon or phosphorus. At each measurement point, an aliquot from the starved culture was diluted into fresh MOPS minimal medium supplemented with 0.2% glucose, distributed in 88-wells of a microtiter plate, and the temporal evolution of optical density (OD_436_) was monitored using a microtiter plate reader. The obtained growth curves were fitted by a constant function and an exponential function, as shown in Fig. 3A. From the fitting, the maximum growth rate *μ* and the lag time (the time at which the constant function and the exponential function intersects in Fig. 3A) was computed for each growth curve. The average specific growth rate and lag time were then computed from the growth curves obtained for each measurement point. Example time courses are shown in Supplementary Fig. S2. We call the lag time obtained in this way the apparent lag time (*λ*). The average growth rate obtained from the fitting is plotted in Supplementary Fig. S3. Also, we have computed the apparent lag time by fitting the data by the growth curve proposed by Baranyi et. al., (Baranyi, 1993). The fitting by Baranyi curve led to approximately the same values of apparent lag time (see section 3 and Fig. S4 in Supplementary Text)

**FIG. 3.**
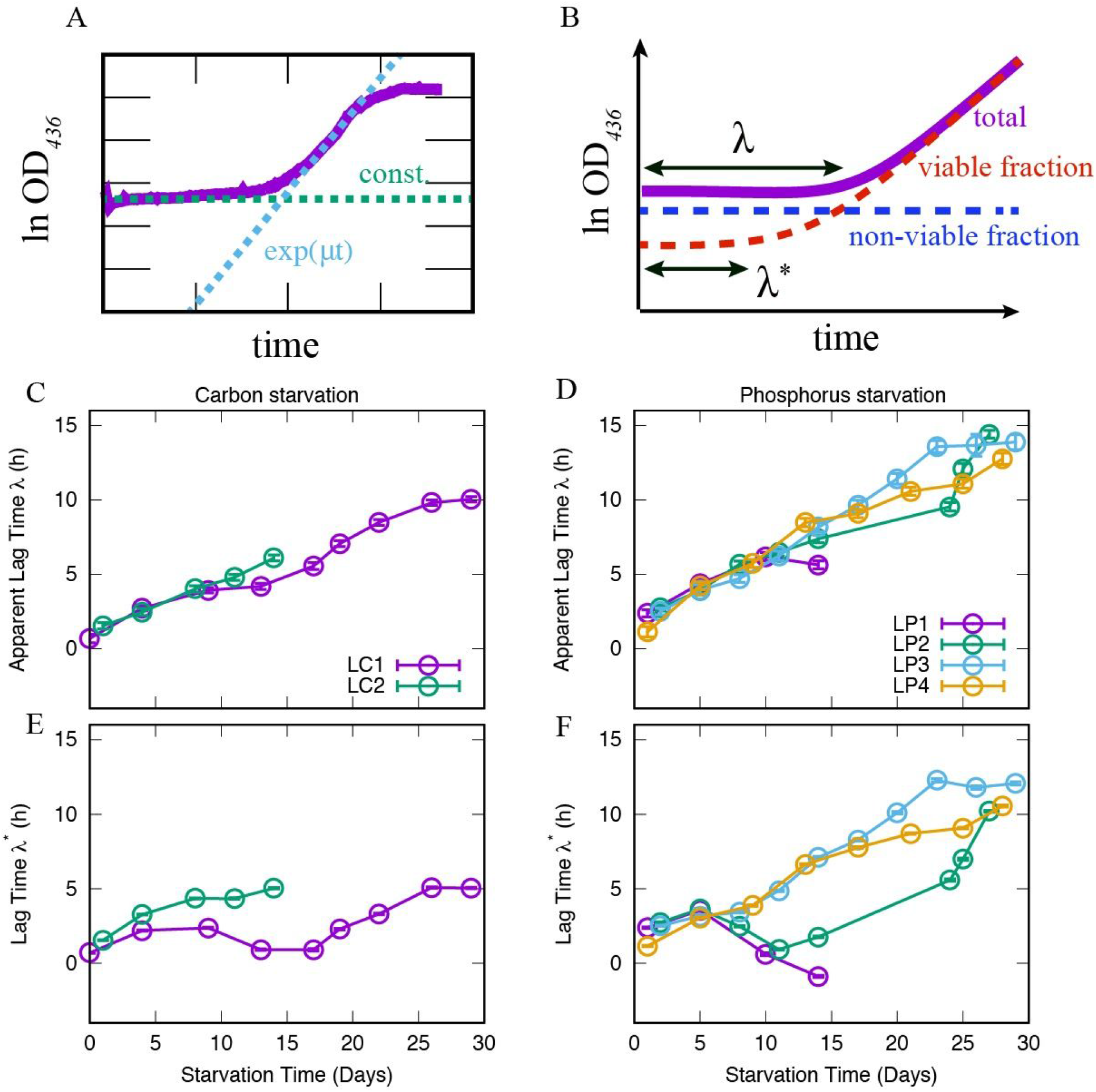
Lag time measurements after carbon or phosphorus starvation. (A). An example of the time series of resuscitation and fitting parameters. The growth curve of the bacterial culture is shown in magenta. The lag phase is fitted by a constant function *f* (dashed green line), and the exponential phase is fitted by an exponential function *g* (dashed blue line). The value of *f*, the slope of *g*, and the intersection point of *f* and *g* corresponds to the initial OD_436_ value (*N*_*0*_), the specific growth rate (*μ*), and the lag time (*λ*), respectively. (B). An illustration of how the effect of non-viable cells is subtracted from the average lag time. We decomposed the total population into viable (red) and non-viable (blue) fractions based on the measured CFU/ml and re-calculated the lag time for the viable fraction (*λ*^*^). (C) The average apparent lag time *λ* for carbon starvation and (D) phosphorus starvation. (E) The lag time *λ*^*\**^ after subtraction of the effect of non-viable cells for carbon starvation and (F) phosphorus starvation. The error bars are the unbiased standard error of the mean (The number of OD curves used to estimate the lag time is typically more than 80. For the propagated errors, see Materials and Methods).

Fig. 3C and FigD show the apparent lag time *λ* as a function of the starvation time for carbon starvation and phosphorus starvation, respectively. Consistent with the literature cited above, the duration of the apparent lag time *λ* increased monotonically with the starvation time for both types of starvation. Furthermore, phosphate starvation had a more pronounced effect on the growth lag than carbon starvation, as cultures starved for phosphate showed a growth lag of several hours already after one day of starvation, while a similar lag time was first observed after ∼4 days of carbon starvation. We also observed a stronger dependence of growth lag on starvation time for the phosphate-starved culture throughout the 30 days of starvation as the slope of the function was steeper for phosphate starvation (0.41 hours/day) than for carbon starvation (0.33 hours/ day).

It should be noted that the measurement of apparent lag time *λ* is affected by the drop in the concentration of viable cells that occurred during starvation because it takes more time for the OD_436_ to increase during re-growth if only a small fraction of the cells that contribute to the initial OD_436_–value can resume growth (see Fig. 3B). Thus, we decomposed the OD_436_ value into two parts, namely the viable fraction and the non-viable fraction based on the value of CFU/ml in the starved culture, the volume of sample transferred to the microtiter plate, and the OD_436_ curves after dilution into fresh growth medium. We further assumed that the OD_436_ of viable cells increased at growth rate *μ* after a certain growth lag from the inoculation time, while the non-viable fraction never resuscitated. Under this assumption, the lag time *λ*^*\**^ subtracted for the death-effect was computed (see section 2 in Supplementary Information).

A schematic description of this calculation is shown in Fig. 3B. The calculated lag time *λ*^*\**^ is shown in Fig. 3E and F for carbon and phosphorus starvation, respectively. The correction for nonviable cells makes the data much noisier. However, the overall correlation between the starvation time and the (corrected) lag time persists. Similarly to the increase of rRNA/cell after day 5 (Fig. 2D), the steep drop of CFU/ml on day 5∼10 in LP1 and LP2 leads to a decrease of the corrected lag time. In addition, the correction even results in negative lag time for LP1. We attribute these problems to the correction method. The subtraction assumes only two types of cells (viable and non-viable fraction) while the important fact that the lag time is distributed among individuals is disregarded. See Supplementary Text section 4 for an extended discussion of this issue.

## Discussion

We have carried out long-term starvation experiments with carbon-limited and phosphate-limited *E. coli* cultures to study the long-term change in the rRNA level, viability, and lag time of bacteria starved for different nutrient types.

Under both starvation conditions, the concentration of full-length rRNA in the culture decreased exponentially with time. The rate of decrease under phosphorus starvation was appreciably higher than that under carbon starvation. This observation is consistent with data from recent short-term starvation experiments (Schink et al., 2019, Fessler et al., 2020), thereby extending the validity of those observations to include rRNA degradation in the late stationary and death phases. Given that RNA is a phosphorus-rich polymer and that most RNA is found in the ribosomes, the more rapid rRNA degradation under phosphorus starvation might be interpreted as cells using the ribosomes as a reservoir for phosphorus. Phosphate starvation leads to a rapid and severe depletion of nucleotide triphosphates, including the cellular energy currency ATP (Kwon et al., 2010). Nucleotides released from rRNA could therefore support essential cellular functions, in particular protein synthesis, enhancing stress survival in the phosphorus-starved cells. On the other hand, the nucleotides stored in rRNA may not benefit carbon-starved cells to the same degree, as the reduction in ATP levels is slow and gradual during carbon starvation (Brauer et al. 2006, Murray et al. 2003). The molecular pathway for rRNA degradation has been discerned (Zundel et al. 2009), but intriguingly, the known regulatory mechanisms do not predict the observed differences in the rates of ribosome degradation under starvation for carbon and phosphorus. Due to the rapid degradation dynamics, we could only quantify the rRNA concentration under phosphorus starvation for up to 16 days, while for carbon starvation we observed that the exponential decrease lasted for 28 days. We refrained from using more sensitive RNA detection methods, such as qRT-PCR, for detection of rRNA late in starvation, because qRT-PCR would not allow us to distinguish full-length rRNA molecules from cleaved degradation products.

The survival kinetics showed distinct behaviors depending on the limiting nutrient. Under carbon starvation, viability decreased exponentially over the starvation period, while cell viability remained approximately constant after an early steep drop (over 5 to 15 days) under phosphorus starvation. Recently, the maintenance energy requirement (Pirt 1965) was shown to set the death rate of cells under carbon starvation (Biselli et al. 2020). Given the similar death kinetics, cell death under carbon starvation is likely led by the same mechanism also in the present experiments. By contrast, the pattern of viability loss under phosphorus starvation is not predicted by the maintenance energy requirement theory. A possible attractive hypothesis to explain the extensive loss of viability a few days into phosphorus starvation is that a certain threshold level of ribosomes per cell is necessary for viability, and that the average bacterium crossed this threshold during the first starvation days.

Far into the starvation period, culturable bacteria under phosphorus starvation showed much lower rRNA levels (rRNA/CFU) than the carbon starvation case. Thus, the carbon-starved cells appear to keep substantially more ribosomes per cell than what is necessary level for viability, while the ribosomes per cell in the phosphorus-starved cells may be close to the critical level.The storage of inactive ribosomes under the carbon-limited condition has been suggested to aid *E. coli* in rapidly accelerating the growth rate during nutrient upshift (Koch 1971, Li et al 2018).

We also measured the re-growth lag time to study if the decreasing rRNA level during the starvation period is reflected in the time it takes bacteria to resume growth upon replenishment of nutrients. The average apparent lag time *λ* showed a linearly increasing trend with the starvation time. The delay in regrowth as cultures have starved for longer could result from an increasing metabolic or regulatory delay in the individual viable cells, a decreasing viable fraction of the biomass that contributes to the cultures’ optical density, or both. After taking into account the effect of the decreasing viability observed in the CFU/ml, the trend became relatively noisy. However, as an overall behavior over about a month, the corrected lag time still showed an increasing trend with increased starvation time. This trend suggests the existence of a metabolic or regulatory delay in individual viable cells that increases during long-term starvation for carbon or phosphorus. Furthermore, the difference in the steepness of the increase in growth lag with increased starvation time implies that differences in the type of nutrient deficiency that induced starvation are persistently reflected in the state of the cells for at least a month, as the amount of rRNA, the duration of the lag time, and the fraction of viable cells stayed qualitatively and quantitatively different between the two starvation conditions. However, we did not observe a clear correlation between the lag time and levels of rRNA per cell for either starvation condition.

We wish to emphasize that all our measurements were carried out at the population level, not at the single-cell level. The calculated value of rRNA per viable cell was relatively constant after the initial drop, but that does not necessarily mean that the amount of rRNA in every individual cell remained constant. As an extreme scenario, the amount of rRNA may decrease over time for some cells, while other cells may even synthesize new rRNA during starvation. Also, for the lag time measurement, there were roughly 10^6^-10^7^ cells in each well of the microtiter plates when the fresh medium was added. As shown by Levin-Reisman et. al (2010) and Moreno-Gámez et al. (2020), the duration of the lag time differs among individuals. If a small fraction of the cells had a relatively short lag time, their offspring would have dominated the whole subculture. In other words, our measurements are likely to reflect how the shortest end of the lag time distribution changes with starvation time.

Phenomenological modeling of bacteria under starved or substrate-limited conditions has also been attracting attention recently (Dai 2017, Himeoka 2017). The model proposed by one of the authors of the present paper (Himeoka 2017) predicts that the lag time increases in a square-root manner with the starvation time. The model prediction fits some experimental data (Pin & Baranyi 2008, Levin-Reisman 2010), but the correspondence is not clear for the data shown in the present paper. There are several scenarios to be tested. Firstly, the population-level measurements could mask important aspects of the lag-time behavior at the single-cell level. Secondly, the functional form of the lag time could depend on the specific setup of the starvation experiment. All of the above-cited starvation experiments were carried out in LB medium, and at low temperatures for one experiment (Pin & Baranyi 2008). These differences could affect the physiological state of the cells leading to distinct results. Further investigation from both the experimental and theoretical sides are needed to fill the gap among those observations and the present result, which could potentially unveil universal features of the bacterial lag phase and stress responses. As far as we know, this study is unique in that we measure the rRNA level and the lag time over a month-long starvation period under well-controlled conditions. Our results demonstrate that even at the endpoint, when the cultures had been starved for one month, carbon-starved cells on average kept a higher level of rRNA per cell than phosphorus-starved cells did. Similarly, after a month of starvation the carbon-starved cells showed shorter lag time than the phosphorus-starved cells, which may support the idea that storage of a ribosome surplus allows carbon-starved cells to respond faster to the reintroduction of nutrient to the medium (Li et al. 2018). All in all, our results suggest that neither cell viability nor lag time after long-term starvation is a simple function of cellular ribosome content. A more stringent examination of this question would require simultaneous measurements of the ribosome level and the lag time at the single-cell level. Such experiments are therefore strongly desired.

## SUPPLEMENTAL MATERIAL

Supplemental material is available online only.

Text S1 including supplemental figures, PDF file.

Table S1, XLSX file.

## ACKNOWLEDGMENTS

This work was supported by the Danish National Research Foundation (DNRF120) and the Independent Research Fund Denmark (8049-00071B and 8021-00280A).

